# Cp36 serine recombinase as a new tool for zebrafish transgenesis

**DOI:** 10.64898/2026.05.06.723361

**Authors:** Savini U. Thrikawala, Bailey Naples, Emily E. Rosowski

## Abstract

One feature key to the versatility of zebrafish as an animal model for biomedical research is the breadth of genetic tools available, including for transgenesis. While the Tol2 transposase system remains the gold standard, its efficiency can be highly variable. Here, we explored the potential of a complementary transgenesis system, Cp36, a large serine recombinase identified from *Clostridium perfringens* previously found to efficiently integrate target cargo into the human genome without a preinstalled attB site. We generated Cp36-based plasmid constructs for zebrafish transgenesis and compared their performance to matched Tol2 plasmids across multiple experimental contexts, including transient expression, germline transmission, and multi-transgene expression. Cp36 integrates small ∼3.5kb cargo into the zebrafish genome and transmits to the next generation as efficiently as Tol2, but Cp36 performance declines substantially for larger ∼7.5kb constructs. Both Cp36 and Tol2 have comparable efficiency in transiently expressing a second construct regardless of the transposase/recombinase used to integrate the first construct, indicating compatibility with sequential transgenesis strategies. In summary, we demonstrate that Cp36 functions as a new alternative transgenesis method in zebrafish.

## Introduction

High fecundity, rapid development, cost-effective maintenance, optical transparency and amenability to genetic manipulation have made zebrafish an indispensable tool in biomedical research. One of the key genetic manipulations that can be performed in zebrafish is transgenesis, which allows for spatio-temporal control of exogenous gene expression, including fluorescent protein markers and genes encoding disease variants. Due to its utility, the toolbox of zebrafish transgenesis has been expanded and optimized for more than 30 years.

Plasmid DNA and bacterial chromosomal injections were first injected into one-cell-stage embryos to generate transgenic zebrafish expressing foreign DNA such as GFP (Stuart, McMurray et al. 1988, Amsterdam, Lin et al. 1995, Jessen, Meng et al. 1998). The foreign DNA integrates randomly via non-homologous recombination, creating mosaic founder zebrafish, albeit with a low germline integration efficiency of around 5-25% (Stuart, McMurray et al. 1988, Culp, Nüsslein-Volhard et al. 1991, Higashijima, Okamoto et al. 1997, Long, Meng et al. 1997). Next, retroviruses were used to deliver DNA into zebrafish embryos. Retroviruses infect cells, reverse-transcribe their RNA genome into DNA, and integrate that DNA into the host genome. This inherent ability was harnessed to deliver and integrate target cargo into the zebrafish genome by injecting a genetically engineered viral vector carrying the transgene (Lin, Gaiano et al. 1994, Gaiano, Allende et al. 1996). This mode of transgenesis created stable insertions at 5-85% germline founder rate, but establishing a productive retroviral vector is experimentally challenging, often non-reproducible, time-consuming, and requires a higher level of biosafety compliance, which limits its use (Amsterdam and Becker 2005, Kawakami 2007).

Plasmids containing Tol2 elements, identified from a naturally-occurring transposon in medaka, are currently the most common technique used for zebrafish transgenesis (Koga, Suzuki et al. 1996, Kawakami, Koga et al. 1998, Kawakami and Shima 1999, Kawakami, Shima et al. 2000, Kemmler, Moran et al. 2023, Hurst, Dimmler et al. 2025). A donor Tol2 plasmid with the cargo sequence is co-injected with in vitro transcribed Tol2 *transposase* mRNA into fertilized zebrafish embryos (Kawakami, Takeda et al. 2004, Urasaki, Morvan et al. 2006, Kawakami 2007). This system employs a “cut-and-paste” mechanism, where the terminal sequences are recognized by the transposase, which cleaves them and then randomly integrates the entrapped cargo sequence into the host genome of somatic and germ cells (Kawakami 2007). This allows random insertion of multiple copies, thereby increasing transgene expression. Injected F0 adults are then outcrossed with wild-type adults, and the F1 embryos are screened to identify F0 founders with successful germline integration of the transgene. An average of 50% to 70% of founders exhibit successful germline integration, but this efficiency can range from as low as 3% to 100% (Urasaki, Morvan et al. 2006, Kawakami 2007). Other transposon systems, the Tc3 transposon isolated from *Caenorhabditis elegans* and the Sleeping Beauty transposon isolated from fish, also have been used in zebrafish but with a low efficiency of germline integration (7% and 31%, respectively) (Raz, van Luenen et al. 1998, Davidson, Balciunas et al. 2003). Therefore, the efficiency of random transgenesis in zebrafish could still be improved.

Large serine recombinases utilize a different mechanism for integrating DNA into the genome. In general, these recombinases recognize a specific attB site in the genome and unidirectionally integrate DNA containing a corresponding attP site (Fogg, Colloms et al. 2014). For example, PhiC31 is a bacteriophage recombinase widely used for targeted transgene integration into genomes of human cells and other experimental models, including zebrafish (Hu, Goll et al. 2011, Mosimann, Puller et al. 2013, Fogg, Colloms et al. 2014, Roberts, Miguel-Escalada et al. 2014, Jusiak, Jagtap et al. 2019, Lalonde, Wells et al. 2024). However, PhiC3 requires a pre-engineered target genome with an attB landing site, which limits both the number of copies of the transgene that can be integrated and the ability to sequentially insert multiple transgenes. Alternatively, some recombinases can use pseudo-attB sites, which are native sequences similar to the attB sequence, as landing sites (Thyagarajan, Olivares et al. 2001). By comparing environmental and clinical bacterial genome isolates, Dunnet *et al*. identified a large serine recombinase, Cp36, from *Clostridium perfringens*, that can target the human genome at multiple locations and is not limited to naturally occurring or engineered attB sites (Durrant, Fanton et al. 2023). In particular, Cp36 recombinase integrated an mCherry cargo of 7.2 kb into human genomes of K562 and HEK293FT cells with 40% efficiency and was able to integrate two genes sequentially (Durrant, Fanton et al. 2023).

Here, we demonstrate that the Cp36 recombinase system can be used for zebrafish transgenesis. Using zebrafish embryos injected with engineered Cp36 donor plasmids and in vitro transcribed Cp36 *recombinase* mRNA, we find that the Cp36 system is equally as efficient as the Tol2 system in integrating cargos of ∼3.5kb, but less efficient for larger ∼7.5kb cargos. We also demonstrate that expression of a second transgene in a previously generated Cp36 transgenic line is possible. Therefore, the Cp36 system can be used as an alternative to the Tol2 system in zebrafish.

## Methods

### Tol2 plasmid constructs

All plasmids generated in this study are listed in Table 1 and were confirmed by Sanger sequencing. Primers used in cloning are listed in Table 2.

**Table 1.** Plasmids generated in this study.

| Plasmid name | Vector backbone | Purpose |
| --- | --- | --- |
| Tol2-*mfap4*:TagRFPT.zf1-GS_cryaa:mCherry | Tol2 | N-terminal tagging vector with expression in macrophages |
| Tol2-*mfap4*:GS-TagRFPT.zf1_cryaa:mCherry | Tol2 | C-terminal tagging vector with expression in macrophages |
| Tol2-*mfap4*:TagRFPT.zf1-GS-atp6v1h_cryaa:mCherry | Tol2 | N-terminal tagged atp6v1h expression in macrophages |
| Tol2-*mfap4*:ctsd-GS-TagRFPT.zf1_cryaa:mCherry | Tol2 | C-terminal tagged ctsd expression in macrophages |
| Tol2-cryaa:mCherry | Tol2 | mCherry expression in the eye |
| Cp36 donor backbone | Cp36 | Base plasmid for insertion of transgenes by Cp36 recombinase |
| Cp36-cryaa:mCherry | Cp36 | mCherry expression in the eye |
| Cp36-*mfap4*:BFP | Cp36 | BFP expression in macrophages |
| Cp36-*mfap4*:TagRFPT.zf1-GS-atp6v1h_cryaa:mCherry | Cp36 | N-terminal tagged atp6v1h expression in macrophages |
| Cp36-*mfap4*:ctsd-GS-TagRFPT.zf1_cryaa:mCherry | Cp36 | C-terminal tagged ctsd expression in macrophages |
| pCS2-Cp36 recombinase-2A-EGFP | pCS2+8 | in vitro transcription of Cp36 *recombinase* mRNA |

**Table 2.**
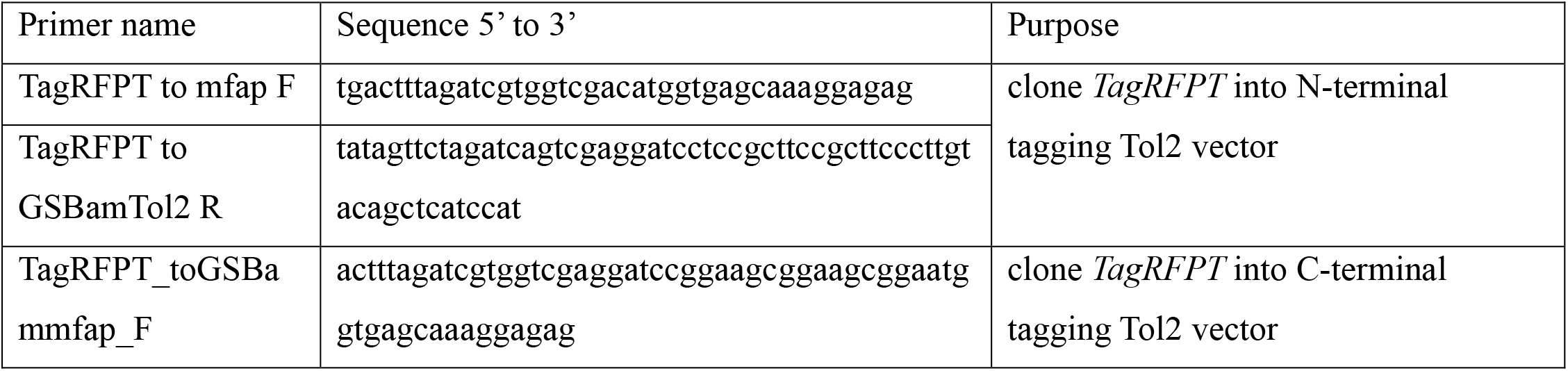
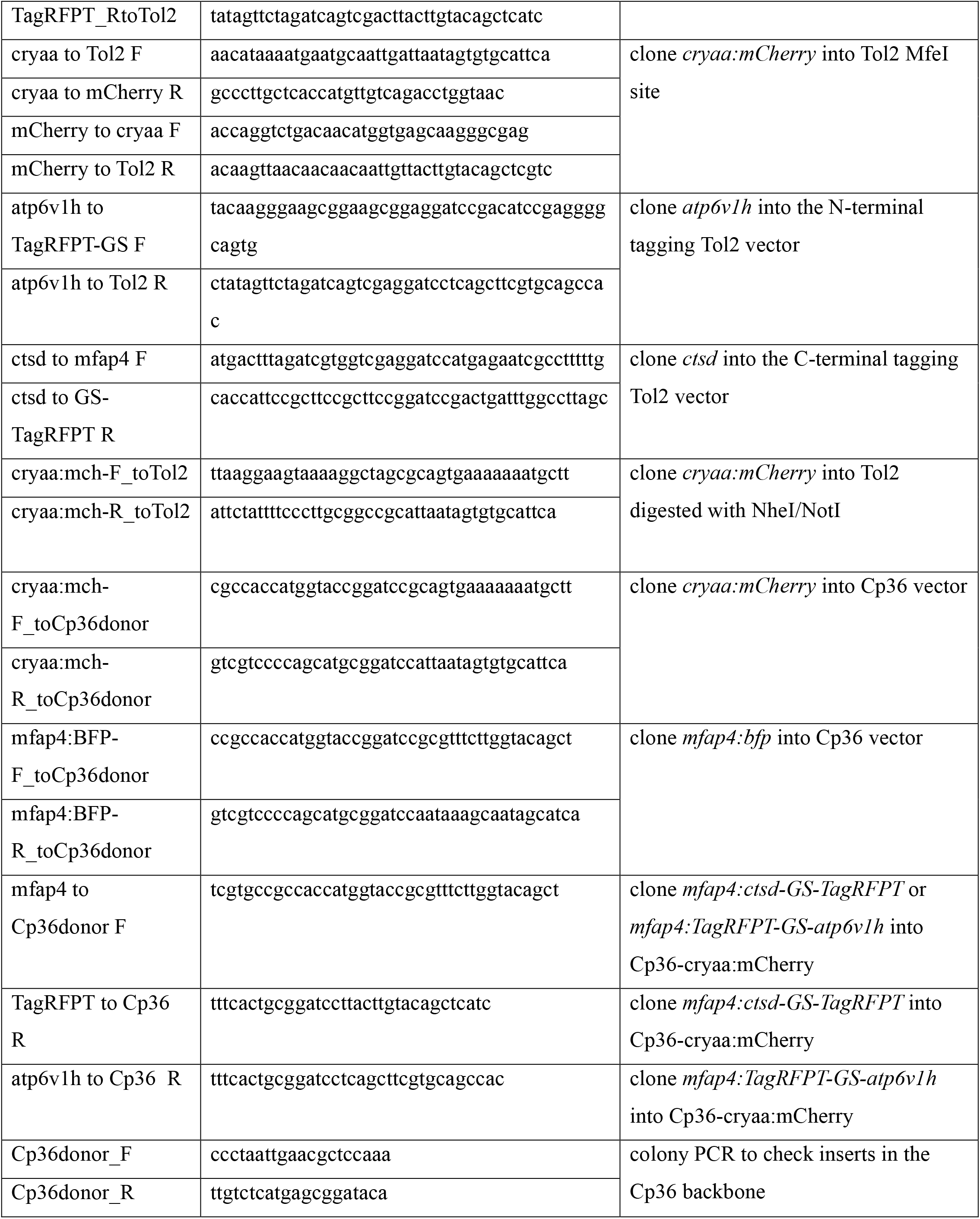
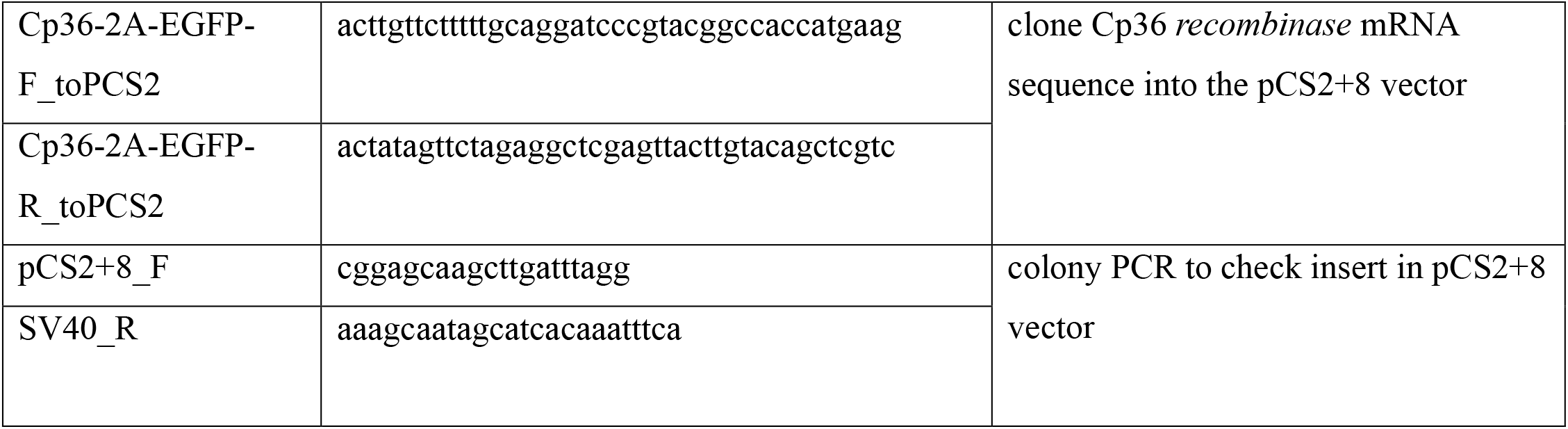
Primer sequences used in the study.

N-terminal and C-terminal tagging vectors under *mfap4* promoter-driven macrophage expression were generated. For the N-terminal vector, a *TagRFPT* sequence codon-optimized for expression in zebrafish amplified from pCS2-TagRFPT.zf1, a gift from Harold Burgess (Addgene plasmid # 61390) (Horstick, Jordan et al. 2015) with Q5 polymerase (NEB) without the stop codon and adding a sequence encoding a 7-amino acid glycine-serine linker, including a BamHI site, at the C-terminal end, was cloned into the SalI site of Tol2-*mfap4* (Addgene plasmid # 232189) (Thrikawala, Anderson et al. 2024) by HiFi cloning (NEB). For the C-terminal vector, the *TagRFPT* sequence was amplified with the stop codon and adding a sequence encoding a 7-amino acid glycine-serine linker, including a BamHI site, at the N-terminal end and cloned into the SalI site of Tol2-*mfap4* by HiFi cloning (NEB). For both of these vectors, a *cryaa* promoter driving mCherry expression in the lens of the eye was also cloned into the MfeI site on the reverse strand by HiFi cloning (NEB), as a marker for genomic integration. Coding sequence for mCherry was amplified from Tol2-*mpx:mCherry-2A-rac2* (a gift from Anna Huttenlocher) (Deng, Yoo et al. 2011) and the sequence of the *cryaa* promoter was amplified from hsp70l-loxP-mCherry-STOP-loxP-H2B-GFP_cryaa:cerulean, a gift from Didier Stainier (Addgene plasmid # 24334) (Hesselson, Anderson et al. 2009). The N-terminal tagging vector was used to generate Tol2-*mfap4:TagRFPT-GS-atp6v1h_cryaa:mCherry* (Supp Fig. 1A), and the C-terminal tagging vector was used to generate Tol2-*mfap4:ctsd-GS-TagRFPT_cryaa:mCherry* (Supp Fig. 1B). Coding sequence of *atp6v1h* and *ctsd* was amplified from cDNA made with iScript RT supermix (Bio-Rad) from total RNA isolated from 2-3 dpf zebrafish larvae with TRIzol (Invitrogen) and cloned into BamHI sites.

Tol2-*cryaa:mCherry* (Supp Fig. 1C) was generated by amplifying *cryaa:mCherry* from the previously generated tagging vectors and cloned by HiFi cloning (NEB) into a Tol2 backbone generated by digesting Tol2-*mfap4* with NheI and NotI to remove the *mfap4* promoter. Tol2-*mfap4:bfp* (Supp Fig. 1D) was previously generated (Thrikawala, Anderson et al. 2024).

### Cp36 donor plasmid constructs

A clean Cp36 donor backbone with an MCS sequence was generated. pAF0185 Cp36_attP-mCherry donor, a gift from Patrick Hsu (Addgene plasmid # 193468) (Durrant, Fanton et al. 2023) was digested with NcoI and SphI (NEB) to remove the mCherry insert, and an oligo containing BamHI and KpnI sites (5’-acggcataatcgtgccgccaccatggtaccggatccgcatgctggggacgacgtcaggtggc-3’) was cloned in by HiFi cloning (NEB).

*cryaa:mCherry* and *mfap4: bfp*, including SV40 polyA sequences, were amplified from Tol2 plasmids described above and cloned into the BamHI site in the Cp36 donor by HiFi cloning (NEB) (Supp Figs. 2A, 2B). *mfap4:TagRFPT*-GS-*atp6v1h* and *mfap4:ctsd*-GS-*TagRFPT*, were amplified from Tol2 plasmids described above and cloned into the KpnI site in Cp36-*cryaa:mcherry* by HiFi cloning (NEB) (Supp Figs. 2C, 2D).

### Generation of Tol2 *transposase* and Cp36 *recombinase* mRNA

A pCS2-Cp36 *recombinase-2A-EGFP* expression plasmid was generated for in vitro transcription of Cp36 *recombinase* mRNA. Cp36 *recombinase-2A-EGFP* was amplified from pAF0084 Ef1a-Cp36-2a-EGFP, a gift from Patrick Hsu (Addgene # 193462) (Durrant, Fanton et al. 2023) and cloned into the pCS2+8 plasmid (a gift from Anna Huttenlocher) digested with BamHI and XhoI (NEB). Cp36 *recombinase* was then in vitro transcribed from NotI-digested (NEB) pCS2-Cp36 *recombinase*-*2A-EGFP*, and Tol2 *transposase* was in vitro transcribed from NotI-digested (NEB) pCS2-*transposase* (a gift from Anna Huttenlocher), using an mMESSAGE mMACHINE SP6 kit (Invitrogen), and mRNA was purified with a MEGAclear kit (Invitrogen).

### Zebrafish lines and maintenance

Adult zebrafish were maintained at 28°C under a 14/10-h light/dark cycle on a centralized filtration life-support system (Aquaneering). All lines were maintained in the wild-type AB background. Embryos were collected after natural spawning and maintained in E3 medium containing methylene blue in petri dishes at 28.5 °C in a 28.5°C incubator.

### Zebrafish microinjections

For Tol2 injections, an injection mix containing 20 ng/μL plasmid, 10 ng/μL Tol2 *transposase* mRNA, cut smart buffer (NEB), and phenol red was used as previously described (Thrikawala, Anderson et al. 2024). To determine the optimum Cp36 plasmid:*recombinase* concentration for transgene expression and to minimize toxicity, different mixes (ng/μL) were prepared (10:10, 10:20, 20:10, 20:20, 30:10, 30:20). For all further Cp36 injections, 20 ng/μL plasmid and 10 ng/μL Cp36 *recombinase* mRNA was used. 1-2 nL of each injection mix was injected into the yolk of single-cell stage embryos. Approximately 6 hours post-fertilization (hpf), unfertilized embryos were removed, and the fertilized embryos were counted, rinsed, and transferred to fresh E3 medium. At 3 dpf, larvae were screened for mCherry or BFP signal using a fluorescent zoomscope (Zeiss SteREO Discovery.V12 PentaFluar with Achromat S 1.0× objective) and the numbers of fluorescent-positive, fluorescent-negative, malformed, and dead larvae were counted.

### Screening for germline integration

At 5-7 dpf, fluorescent positive F0 larvae were transferred to adult housing tanks (∼50 larvae per tank) and raised to adulthood. Around 3 months post-fertilization (mpf), the number of F0 adults surviving in each housing tank and their sex were counted. These F0 adults were then outcrossed individually with wild-type adults, and the F1 embryos were collected and screened for fluorescent expression at 3 dpf as described above. At least 20 larvae per F0 adult were screened. In case of lower embryo production, the F0 adult was spawned again to screen more larvae. The total number of F1 larvae and the number of fluorescent-positive F1 larvae were counted.

## Results

### Cp36 recombinase system can mediate transient mosaic expression in F0 zebrafish larvae

The Tol2 transposase system has long been used by us and others to generate stable zebrafish transgenic lines (Kawakami 2007, Kemmler, Moran et al. 2023, Thrikawala, Anderson et al. 2024, Hurst, Dimmler et al. 2025). Here, we investigate the potential of utilizing a distinct recombinase, Cp36, which efficiently integrates transgenes into the human genome (Durrant, Fanton et al. 2023), for zebrafish transgenesis. We generated a plasmid vector containing the Cp36 attP site, which is recognized by Cp36 recombinase to integrate the vector into the target zebrafish genome, and an mCherry coding sequence under the eye lens-specific zebrafish *cryaa* promoter (Cp36-*cryaa:mCherry*) to visually identify larvae with transgene expression (Fig. 1A, Supp Fig. 2A). We then generated a pCS2 plasmid containing the Cp36 recombinase coding sequence, which was then used to transcribe the mRNA in vitro. We first optimized the concentrations of the Cp36_attP vector and *recombinase* mRNA to be microinjected. We injected a range of different vector:*recombinase* mRNA mixtures and fluorescently screened the F0 larvae at 3 dpf for mCherry expression in the eye (Fig. 1B), to find that the final vector:*recombinase* mRNA concentrations of 20:10 ng/μL were the most efficient (Fig. 1C). These are the same concentrations we have previously used for Tol2 vectors and *transposase* mRNA (Thrikawala, Anderson et al. 2024).

**Figure 1.**
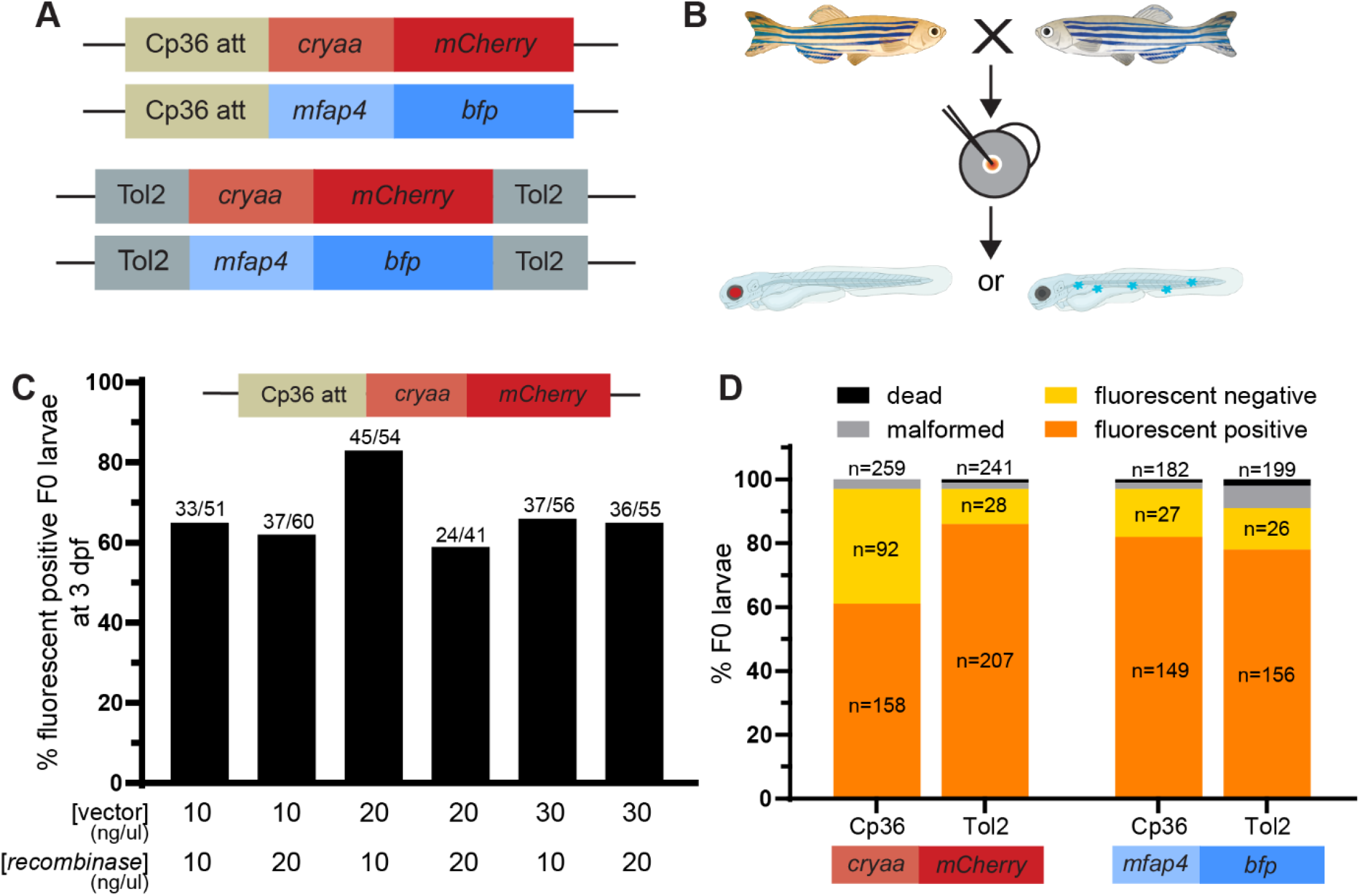
Transient transgene expression with Cp36 recombinase system. (A) Schematic of the four plasmid constructs cloned and tested. A *cryaa* promoter driving mCherry and an *mfap4* promoter driving BFP were cloned after the Cp36 att site or between Tol2 arms. (B) Wild-type adults were spawned and the plasmid constructs together with their respective *recombinase/transposase* mRNA were injected into 1 cell stage embryos. F0 larvae were screened for fluorescence expression at 3 dpf. (C) Embryos were injected with different concentration combinations of the Cp36-*cryaa:mCherry* plasmid and Cp36 *recombinase* mRNA to find optimal concentrations. (D) Wild-type embryos were injected with each construct and the respective *recombinase/transposase* mRNA. At 3 dpf, the number of fluorescent positive, fluorescent negative, malformed, and dead F0 larvae were counted. The total larval n for each category is indicated.

We next compared the efficiency of Cp36 recombinase with Tol2 transposase. We generated a corresponding Tol2 vector to express mCherry under the *cryaa* promoter (Tol2-*cryaa:mCherry*) (Fig. 1A, Supp Fig. 1C). We also expanded our comparison to a different zebrafish promoter, *mfap4*, and generated a Cp36_attP vector to express BFP under this promoter (Cp36-*mfap4:bfp*) (Fig. 1A, Supp Fig. 2B). The equivalent Tol2-*mfap4:bfp* vector was previously generated (Fig. 1A, Supp Fig. 1D) (Thrikawala, Anderson et al. 2024). We microinjected all four vector constructs into wild-type embryos with the corresponding *transposase*/*recombinase* mRNA, and screened 3 dpf F0 larvae for transient fluorescent expression (Fig. 1B). Compared to Tol2, Cp36 is slightly lower in efficiency for expression of cryaa:mCherry, while both Tol2 and Cp36 show similar efficiency in expressing BFP in macrophages in F0 larvae (Fig. 1D). We also quantified the percentage of malformed embryos, which was lower with Cp36 injections as compared to Tol2 injections, but lower than 7% in both systems indicating minimal toxicity from the injections (Fig. 1D). Overall, both Cp36 and Tol2 have similar efficiency in transiently expressing fluorescent cargo in F0 larvae.

### Cp36 has a comparable efficiency to Tol2 in the germline transmission of transgenes

The fluorescent expression in F0 larvae does not necessarily demonstrate integration of the plasmid construct into the genome as transient expression from plasmids can also occur. We therefore raised all fluorescent-positive F0 larvae to adulthood to determine the efficiency of germline integration. First, we tested the effect of plasmid injections on long-term survival by monitoring the number of F0 individuals reaching adulthood. While individuals injected with the Cp36-*cryaa:mCherry* construct had a lower survival than their Tol2 counterparts, for the *mfap4:bfp* constructs, Cp36 and Tol2 have similar survival (Fig. 2A), indicating embryos injected with Cp36 plasmids can survive to adulthood. To test germline integration, we outcrossed F0 adults with wild-type adults and screened F1 larvae for fluorescence expression to determine if the transgene is transmitted to their progeny. At least 20 larval progeny were screened per F0 individual to assess successful germline transmission. For Cp36 constructs, we found ∼12% of F0 founders transmit the *cryaa:mCherry* construct and ∼8.5% transmit the *mfap4:bfp* construct, demonstrating that this system can mediate germline integration of transgenes (Fig. 2B). However, these percentages were lower than the matched Tol2 constructs which transmitted in ∼57% and ∼19% of F0s, respectively (Fig. 2B). For the F0 adults that do transmit the transgenes, we also quantified what percentage of their progeny were fluorescent positive. The Cp36*-cryaa:mCherry*-injected F0 adults give rise to an average of 8% fluorescent progeny as compared to 18% by Tol2*-cryaa:mCherry*-injected F0 adults (Fig. 2C). For *mfap4:bfp* constructs, the average rates of germline transmission to F1 larvae are similar for Cp36 and Tol2, at 13% and 11%, respectively (Fig. 2C). Overall, Cp36 recombinase can mediate germline integration of transgenes, however in some cases the efficiency of this integration may be slightly lower than Tol2 transposase.

**Figure 2.**
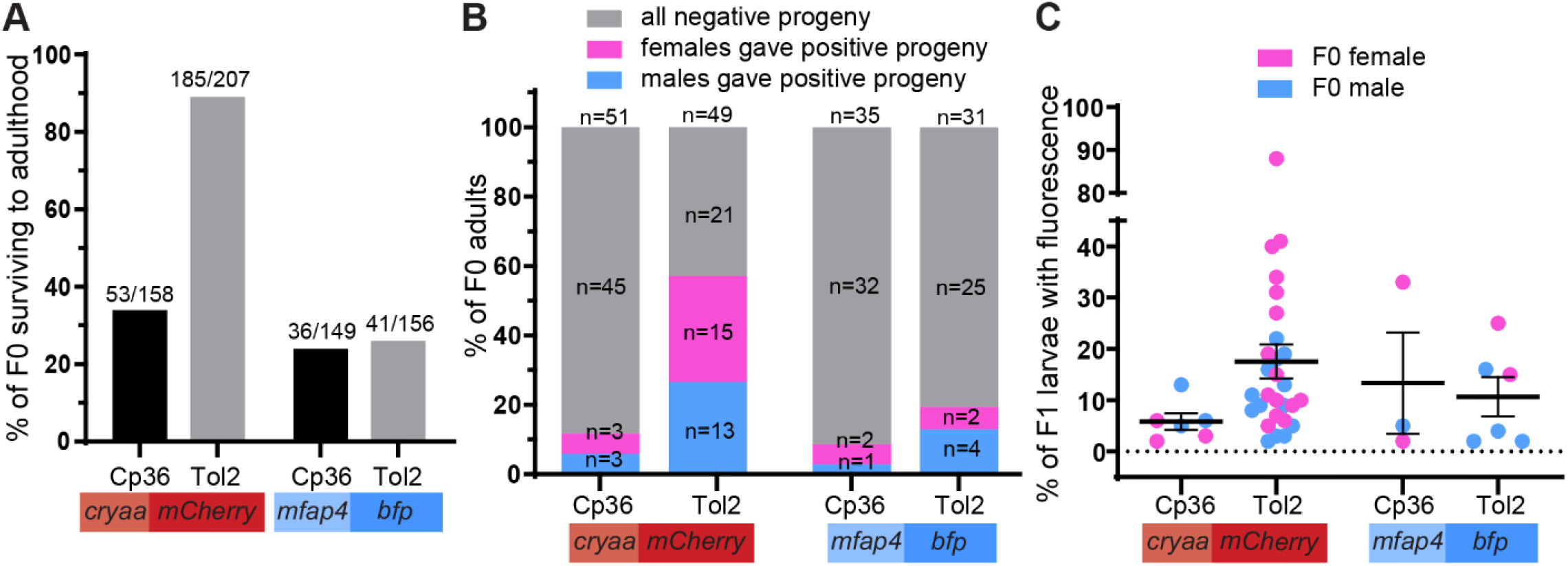
Cp36 recombinase can mediate germline integration of transgenes. (A) F0 larvae positive for transgene expression were raised and the number surviving to adulthood was counted at ∼2 months. (B) F0 adults were outcrossed with wild-type and the resulting F1 larvae were screened for transgene expression. The total adult n screened for each transgene is indicated. (C) For the F0 adults that gave fluorescent positive F1 progeny, the percentage of fluorescent positive progeny was quantified. Each data point represents an individual F0 adult, color-coded by sex, and mean ± SEM across all F0 adults is indicated.

### Cp36 is less efficient in integrating and transmitting larger transgenes

Both the *cryaa:mCherry* and *mfap4:bfp* transgene cargos are ∼3.5 kb. However, Tol2 can efficiently integrate cargo of >11 kb (Urasaki, Morvan et al. 2006). We therefore next asked if Cp36 can integrate larger cargos. We generated both Cp36 and Tol2 plasmids expressing a zebrafish gene (either *atp6v1h* or *ctsd*) tagged with a fluorescent protein TagRFPT under the macrophage-specific *mfap4* promoter. These plasmids also contain the *cryaa:mCherry* eye marker to facilitate screening of larvae, making them ∼7.5 kb total (Fig. 3A, Supp Figs. 1, 2). We injected wild-type embryos with these four new constructs and their respective *recombinase*/*transposase* mRNA and screened F0 larvae for the eye marker. We found ∼50% F0 larvae injected with the Cp36 constructs and ∼80% of Tol2 constructs expressed the mCherry eye markers, indicating successful delivery of the cargo, with low levels of dead or malformed larvae (Fig. 3B). 30-50% of these larvae survived to adulthood, also suggesting no toxicity associated with plasmid injections (Fig. 3C). F0 adults were outcrossed with wild-type adults to identify individuals with germline integration of the cargo. Across all constructs, the majority of these F0 adults produce fluorescent negative progeny, indicating no germline integration (Fig. 3D). For both Cp36 gene constructs, only 7% of the F0 adults have germline integration (Fig. 3D). In contrast, ∼40% of F0 adults njected with Tol2 constructs have germline integration of the larger cargo (Fig. 3D), suggesting that the germline integration efficiency is independent of the transgene, but dependent on the transposase/recombinase system used. For the adults that transmit transgenes, we then quantified the percentage of their progeny that were fluorescent positive. For both gene constructs, Tol2 produces a higher percentage of fluorescent positive F1 larvae than Cp36 (Fig. 3E). Overall, our results indicate that Cp36 is comparable in efficiency to Tol2 in integrating and transmitting smaller cargo (∼3.5 kb), but not larger cargo (∼7.5 kb) into the zebrafish genome.

**Figure 3.**
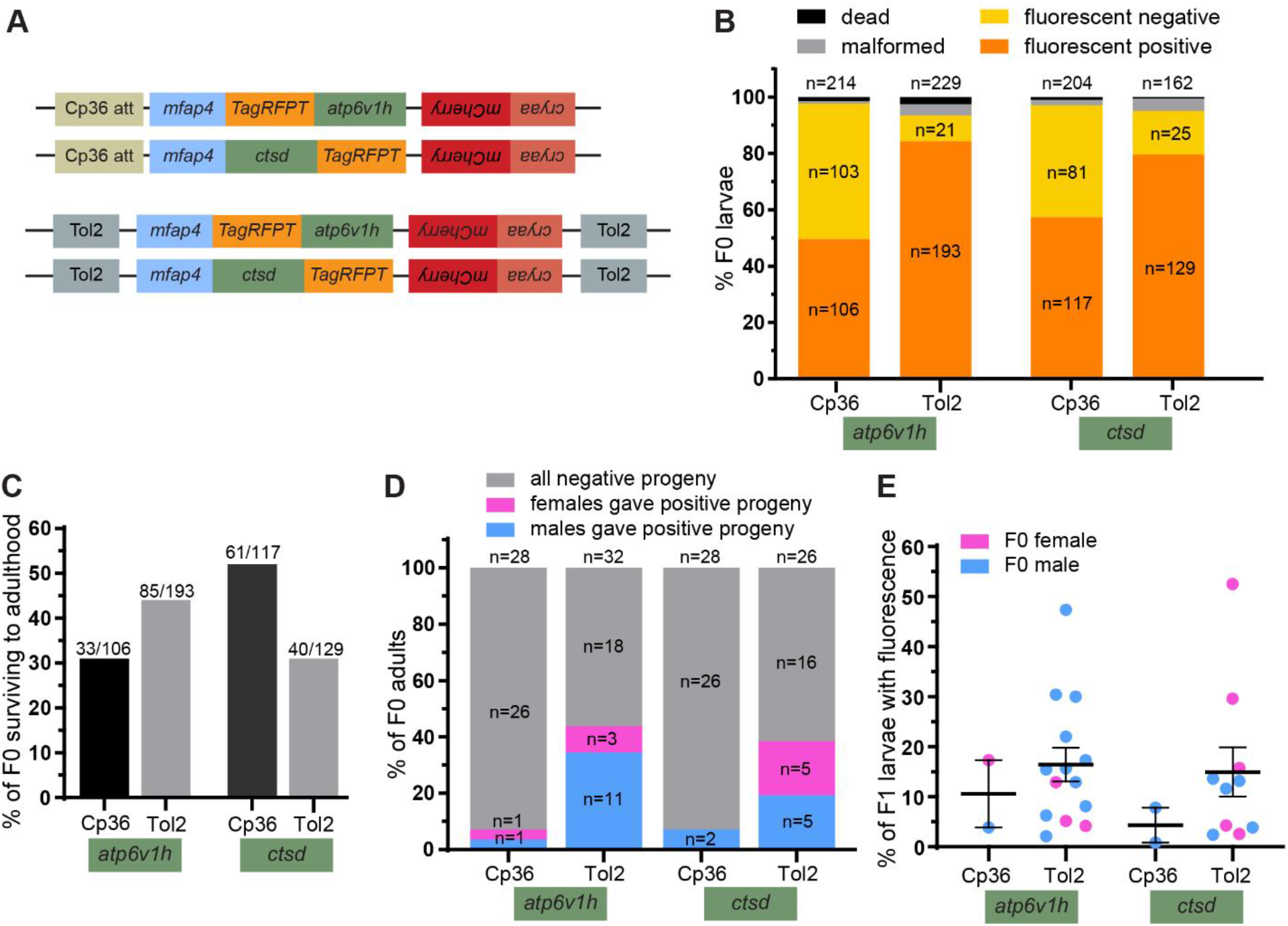
Cp36 recombinase is less efficient than Tol2 transposase in germline integration of larger transgenes. (A) Schematic of two different large constructs (∼7.5 kb) cloned under the Cp36 att site or between Tol2 arms. (B) Wild-type adults were spawned and the plasmid constructs together with their respective *recombinase/transposase* mRNA were injected into 1 cell stage embryos. F0 larvae were screened for fluorescent expression at 3 dpf. The number of fluorescent positive, fluorescent negative, malformed, and dead were counted. The total larval n for each category is indicated. (C) F0 larvae positive for transgene expression were raised and the number surviving to adulthood was counted at ∼2 months. (D) F0 adults were outcrossed with wild-type and the resulting F1 larvae were screened for transgene expression. The total adult n screened for each transgene is indicated. (E) For the F0 adults that gave fluorescent positive F1 progeny, the percentage of fluorescent positive progeny was quantified. Each data point represents an individual F0 adult, color-coded by sex, and mean ± SEM across all F0 adults is indicated.

### Cp36 can deliver a second fluorescent cargo into larvae that already express a fluorescent transgene

Another important consideration for a transgenesis system is the ability to express a second transgene in an already transgenic background. Cp36 can transmit a second construct into human cells without losing the first cargo (Durrant, Fanton et al. 2023), and we tested whether this is also possible in zebrafish. We outcrossed Cp36-*cryaa:mCherry* and Tol2*-cryaa:mCherry* stable transgenic F1 adults with wild-type adults and injected the embryos with Cp36- or Tol2-*mfap4:bfp* constructs and the respective *recombinase* or *transposase* mRNA (Fig. 4A). All resulting larvae were first screened for mCherry expression to select larvae with the previous stable transgene. mCherry positive larvae were then screened for BFP expression, which would indicate at least transient expression of the second cargo. From the progeny of Cp36 F1 adults, we find 39% of mCherry positive larvae also express the second BFP Cp36 construct (Fig. 4B, first bar), demonstrating that a second Cp36 cargo can be expressed in larva already carrying a stable Cp36 transgene. This rate is similar for the progeny of Tol2 F1 adults expressing a second Tol2 cargo, with 33% of Tol2 mCherry positive larvae also expressing a BFP Tol2 construct (Fig. 4B, last bar). Therefore, Cp36 can be utilized for successive transgenesis with the same efficiency as Tol2.

**Figure 4.**
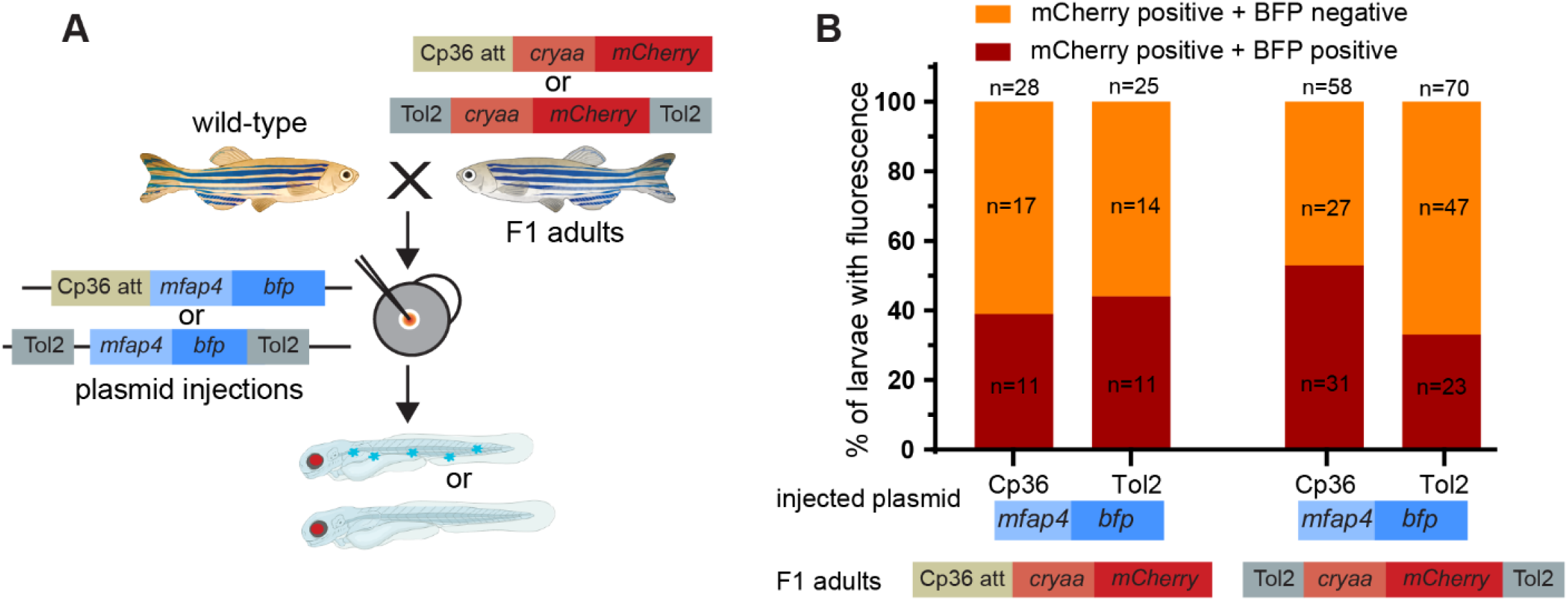
Cp36 recombinase can mediate expression of a second transgene. (A) F1 adults with Cp36-*cryaa:mCherry* or Tol2-*cryaa:mCherry* transgenes were outcrossed with wild-type adults and the embryos were injected with Cp36-*mfap4:bfp* or Tol2-*mfap4:bfp* plasmids and their respective *recombinase/transposase* mRNA. Larvae were screened for fluorescent transgene expression at 3 dpf. (B) Of mCherry positive larvae, the percentage expressing BFP and not expressing BFP was quantified. The total larval n for each category is indicated.

We also aimed to test cross-compatibility between these systems. From the progeny of Cp36 F1 adults, we find 44% of mCherry positive larvae also express a second BFP Tol2 construct (Fig. 4B, second bar). Similarly, from the progeny of Tol2 F1 adults, we find 53% of mCherry positive larvae also express a second BFP Cp36 construct (Fig. 4B, third bar). There is therefore a slight increase in the percentage of larvae expressing a second transgene when a different system is used (44% and 53% positivity versus 39% and 33%). Overall, both Cp36 and Tol2 can deliver a second fluorescent cargo into zebrafish larvae that already express another fluorescent cargo, regardless of the transposase/recombinase used to deliver this first cargo, demonstrating that Cp36 is also amenable for sequential transgenesis in zebrafish.

## Discussion

The development of efficient and flexible transgenesis tools remains central to advancing zebrafish as a model for vertebrate biology and disease research. In this study, we evaluated the potential of the Cp36 large serine recombinase system as an alternative to the widely used Tol2 transposase system for generating transgenic zebrafish. Our findings demonstrate that Cp36 can mediate genomic integration and germline transmission of transgenes in zebrafish, comparable to Tol2, establishing it as a functional addition to the zebrafish transgenesis toolkit.

A major limitation of the Cp36 system revealed in this study is the reduced efficiency in integrating larger transgenes (∼7.5kb). While Tol2 maintained relatively high germline transmission rates for these larger constructs, Cp36 showed a marked decrease in both founder frequency and transmission efficiency. This size-dependent limitation suggests that Cp36-mediated recombination may be less tolerant of larger cargo, potentially due to constraints in recombinase activity, DNA processing, or integration stability. It should be noted that, unlike Tol2, which underwent multiple rounds of optimization (Kawakami, Koga et al. 1998, Kawakami and Shima 1999, Kawakami, Shima et al. 2000, Kawakami, Takeda et al. 2004, Urasaki, Morvan et al. 2006, Kawakami 2007, Kawakami, Asakawa et al. 2016), the Cp36 recombinase we used consisted of unaltered bacterial sequences. While our data shows promise, Cp36 may be able to be further improved to increase efficiency. For example, we did not codon-optimize the recombinase sequence for zebrafish.

Large recombinases identify an attP landing site to integrate a single copy of the transgene (Fogg, Colloms et al. 2014). Such recombinases have successfully been used in zebrafish previously, primarily with the PhiC31 integrase (Mosimann, Puller et al. 2013, Lalonde, Wells et al. 2024). In this pIGLET system, attP landing sites were integrated into the zebrafish genome in single copies using Tol2 transgenesis. This site can then efficiently integrate plasmids containing transgenes and the corresponding attB site(Mosimann, Puller et al. 2013, Lalonde, ells et al. 2024). The pIGLET system has several advantages, including a known single insertion site that leads to reproducible expression, both between transgenes and across generations(Lalonde, Wells et al. 2024). However, there are some drawbacks to this system. First, some cell-type specific promoters, including the macrophage-specific promoters *mpeg1* and *mfap4* are relatively weak and require multi-copy integration for detectable fluorescence expression (Ellett, Pase et al. 2011, Walton, Cronan et al. 2015). Second, certain experimental setups may require the use of zebrafish with two or three transgenes integrated into the same genome. We therefore focused on a recombinase that was found to integrate multiple copies of the target cargo into the human genome at pseudo-attB sites, sequences that are similar to a consensus Cp36 attB sequence, but that are found naturally in the human genome (Durrant, Fanton et al. 2023). We demonstrate that this Cp36 system can also stably integrate transgenes into the zebrafish genome, but it is still unknown if this integration also targets pseudo-att sites or how many copies of the transgene have been integrated. Importantly, our results also demonstrate that Cp36 can introduce a second transgene into embryos already carrying an existing fluorescent reporter, regardless of whether the initial transgene was delivered by Cp36 or Tol2. Interestingly, we observe a reciprocal efficiency pattern: Tol2 performs better when Cp36 is used for the first integration, and Cp36 performs better when Tol2 is used initially. This may be because these two systems have different preferences for genomic locations of integration.

Overall, our study establishes Cp36 recombinase as a functional system for mediating transgene integration and germline transmission in zebrafish. While its efficiency is comparable to Tol2 for smaller constructs, its reduced performance with larger cargo highlights the need for further optimization. Cp36 is a versatile tool for zebrafish transgenesis because it can be used for random integration of multiple transgene copies and sequential transgenesis. Together, these findings establish Cp36 as a viable, mechanistically distinct alternative to transposon-based systems, expanding the zebrafish transgenesis toolkit.

## Data availability statement

The data underlying this article are available in the article and in its online supplementary material.

## Acknowledgements

We thank Scott Horman and the Aquatic Animal Research Laboratory for assistance with fish system aintenance and for providing the microinjection set-up for embryo injections. We thank all the members of the Rosowski Lab for helpful discussions and for assistance with zebrafish care.

