## Supplemental Figs 1 and 2 for "Cp36 serine recombinase as a new tool for zebrafish transgenesis"

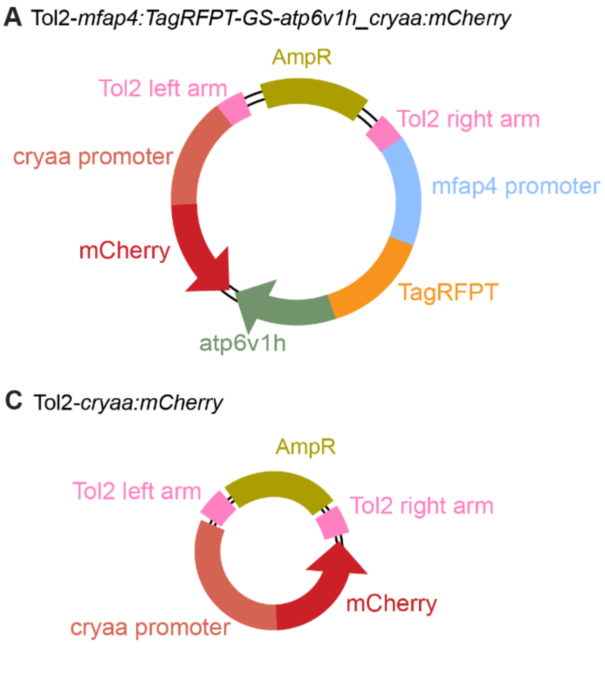


**Supplemental Figure 1.** Plasmids generated in the Tol2 backbone. (A, B) N-terminal and C-terminal TagRFPT tagging vectors under *mfap4:*driven macrophage expression were generated. A *cyraa* promoter driving mCherry was also cloned on the reverse strand. (A) N-terminal tagging vector was used to generate Tol2-*mfap4:TagRFPT-GS-atp6v1h_cryaa:mCherry* by cloning PCR-amplified *atp6v1h*. (B) C-terminal tagging vector was used to generate *mfap4:ctsd-GS-TagRFPT* by cloning PCR-amplified *ctsd*. (C) Tol2-*cryaa:mCherry* was generated by amplifying *cryaa:mCherry* from the previous plasmids and cloning into a Tol2 backbone generated by digesting Tol2-*mfap4* with NheI and NotI to remove the *mfap4* promoter. (D) Tol2-*mfap4:BFP* was previously generated (Thrikawala, Anderson et al. 2024).

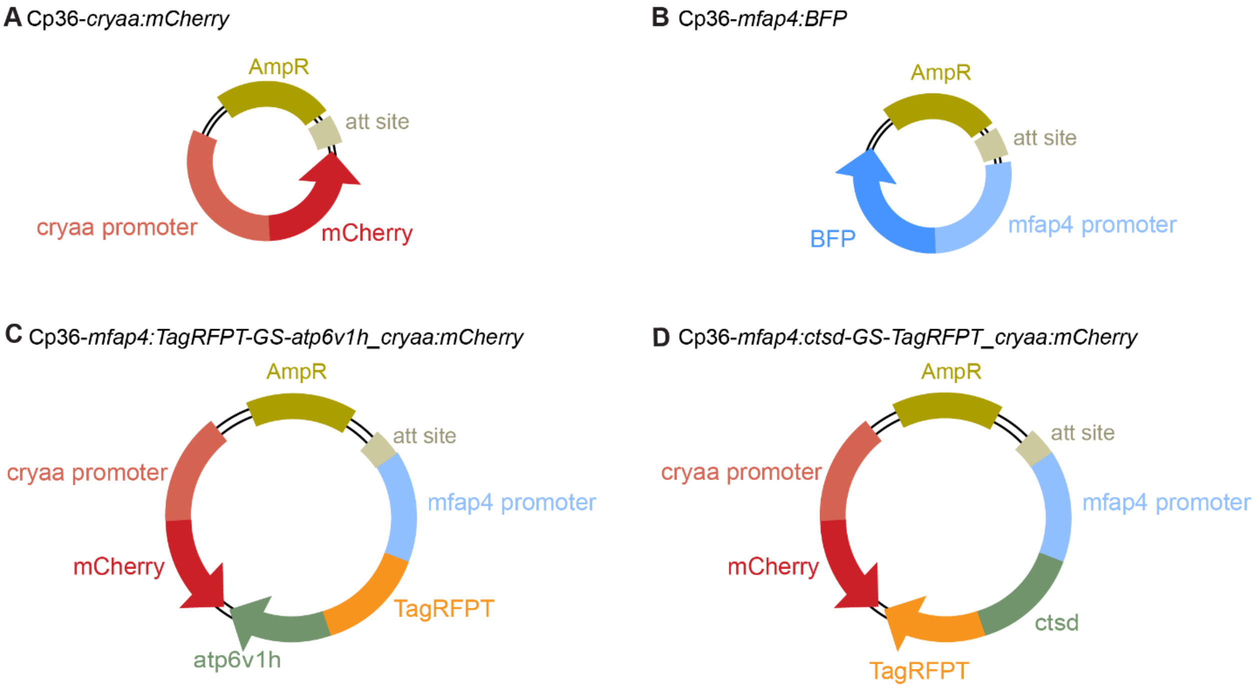


**Supplemental Figure 2.** Plasmids generated in the Cp36 backbone. A clean Cp36 donor backbone with an MCS sequence was generated. (A, B) *cryaa:mCherry* and *mfap4:BFP* were amplified from the plasmids shown in Supp Fig. 1 and cloned into the Cp36 backbone. (C, D) *mfap4:TagRFPT*-GS-*atp6v1h* and *mfap4:ctsd*-GS-*TagRFPT* were amplified from the plasmids shown in Supp Fig. 1 and cloned into the Cp36 backbone.
